# reguloGPT: Harnessing GPT for Knowledge Graph Construction of Molecular Regulatory Pathways

**DOI:** 10.1101/2024.01.27.577521

**Authors:** Xidong Wu, Yiming Zeng, Arun Das, Sumin Jo, Tinghe Zhang, Parth Patel, Jianqiu Zhang, Shou-Jiang Gao, Dexter Pratt, Yu-Chiao Chiu, Yufei Huang

**Affiliations:** Electrical and Computer Engineering, University of Pittsburgh; Hillman Cancer Center, University of Pittsburgh Medical Center; Division of Hematology/Oncology, Department of Medicine, University of Pittsburgh; Department of Electrical and Computer Engineering, The University of Texas at San Antonio; Department of Microbiology and Molecular Genetics, University of Pittsburgh; School of Medicine, UC San Diego

**Keywords:** Molecular Regulatory Pathways, Knowledge Graph, GPT, In Context Learning, m^6^A mRNA Methylation

## Abstract

**Motivation:** Molecular Regulatory Pathways (MRPs) are crucial for understanding biological functions. Knowledge Graphs (KGs) have become vital in organizing and analyzing MRPs, providing structured representations of complex biological interactions. Current tools for mining KGs from biomedical literature are inadequate in capturing complex, hierarchical relationships and contextual information about MRPs. Large Language Models (LLMs) like GPT-4 offer a promising solution, with advanced capabilities to decipher the intricate nuances of language. However, their potential for end-to-end KG construction, particularly for MRPs, remains largely unexplored.

**Results:** We present reguloGPT, a novel GPT-4 based in-context learning prompt, designed for the end-to-end joint name entity recognition, N-ary relationship extraction, and context predictions from a sentence that describes regulatory interactions with MRPs. Our reguloGPT approach introduces a context-aware relational graph that effectively embodies the hierarchical structure of MRPs and resolves semantic inconsistencies by embedding context directly within relational edges. We created a benchmark dataset including 400 annotated PubMed titles on N6-methyladenosine (m^6^A) regulations. Rigorous evaluation of reguloGPT on the benchmark dataset demonstrated marked improvement over existing algorithms. We further developed a novel G-Eval scheme, leveraging GPT-4 for annotation-free performance evaluation and demonstrated its agreement with traditional annotation-based evaluations. Utilizing reguloGPT predictions on m^6^A-related titles, we constructed the m^6^A-KG and demonstrated its utility in elucidating m^6^A’s regulatory mechanisms in cancer phenotypes across various cancers. These results underscore reguloGPT’s transformative potential for extracting biological knowledge from the literature.

**Availability and implementation:** The source code of reguloGPT, the m^6^A title and benchmark datasets, and m^6^A-KG are available at: https://github.com/Huang-AI4Medicine-Lab/reguloGPT.

## I. Introduction

Molecular Regulatory Pathways (MRPs) are central to our understanding of biological functions, as they reveal how genetic variations or chemical stimuli influence biological processes and diseases. Studying MRPs allows scientists to uncover the molecular mechanisms controlling biological functions, aiding in the identification of disease-contributing dysregulations and guiding the development of targeted therapies. As such, elucidating MRPs is a key goal in biomedical research, offering critical insights into biological processes and informing the design of precise medical treatments. For organizing and analyzing the extensive data within MRPs, Knowledge Graphs (KGs) have become instrumental. These KGs offer structured representations of complex biological knowledge, detailing the interactions among various entities such as genes, proteins, and biological processes [1], [2]. While databases like KEGG, Reactome, and QIAGEN Ingenuity Pathway Analysis have been established through meticulous human curation, the sheer volume and pace of new research publications pose a significant challenge to these manual efforts. To address this, automated Natural Language Processing (NLP) methods have been developed, combining rule-based and machine-learning strategies to improve the extraction of biomedical knowledge from literature, as seen in databases like RepoDB, MSI, Hetionet, DrugMechDB, and INDRA [3].

Current tools for mining gene associations are inadequate for mapping complex MRPs, which involve intricate relationships and hierarchical structures. For example, the sentence “METTL3-mediated m^6^A methylation of SPHK2 promotes gastric cancer progression by targeting KLF2.” suggests a context-specific graph of N-ary relationships involving several entities, i.e., METTL3, m^6^A (N6-methyladenosine), SPHK2, KLF2, progression and gastric cancer as the context (Fig. 1). This graph encompasses both explicit and implicit regulatory relationships that collectively describe the mechanism by which METTL3 regulates the progression of gastric cancer. Extracting such detailed graphs from MRP descriptions challenges existing NLP methods and requires advanced Named Entity Recognition (NER) and N-ary Relationship Extraction (RE), along with context identification.

**Fig. 1.**
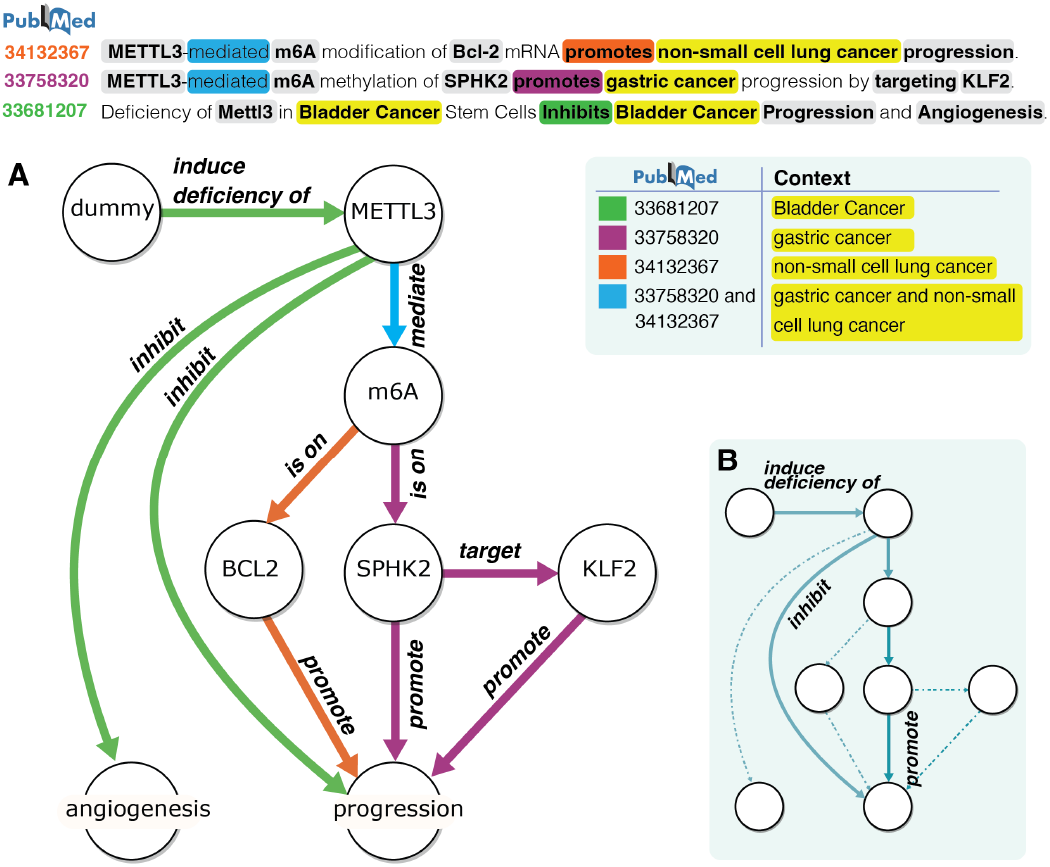
(A) reguloGPT builds a context-aware knowledge graph (KG) based on PubMed sentences depicting molecular regulatory pathways. The KG reflects the hierarchy of molecular pathways, while also incorporating extracted regulatory contexts and associated PubMed IDs into edges. This enables the delineation of context-specific regulation. (B) The exclusion of context in KG could introduce contradictory relations or wrong conclusions in the downstream pathway. For example, the highlighted path suggests the ‘inhibition’ and ‘promotion’ of ‘progression’ with ‘induced deficiency of’ METTL3, which is incorrect.

Existing NLP methods for biomedical KG construction can be categorized as rule-based including SemRep [4] and REACH [5], machine-learning based including EIDOS [6] and GNBR [7], or a mix of the two such as Turku Event Extraction System (TEES) [8]. However, they focus on binary relationships, represented as triplets (entity A, relationship r, entity B) [9], and struggle with N-ary relationships in MRPs. This limitation leads to cascading errors from misidentified entities to RE and redundant/missed relationships, and increased complexity [10]. Specifically, REACH [5] uses a rule-based approach to effectively identify entities and relationships within biomedical texts. Similarly, EIDOS [6] is tailored for extracting structured information from scientific literature, employing machine learning techniques to recognize entities and relationships, thus boosting its information extraction capabilities. TEES [8], aims to extract events and participants from biomedical texts, combining rule-based methods with machine learning. In a similar vein, GNBR [7] specializes in normalizing gene mentions and extracting binary relations from biomedical literature. GNBR employs machine learning for effective gene-related information extraction. SemRep [4] is another system that utilizes a rule-based methodology for biomedical relation extraction. It is designed to navigate and interpret complex language structures in biomedical literature, focusing on the extraction of meaningful semantic relations.

N-ary relationships, involving more than two entities, are crucial for a comprehensive representation of biological interactions. While N-ary RE is well-investigated in general KG construction [11], it is under-explored in the biomedical domain, although a few recent works have considered the prediction of drug-gene-mutation relationships and others from multiple sentences [12]. Furthermore, existing methods are limited in capturing important contextual information like diseases and tissue types, potentially leading to inconsistencies in and misinterpretation of biomedical KGs.

The advent of Large Language Models (LLMs) like GPT-4 represents a significant leap forward in NLP, providing deep insights into the contextual dynamics of language. These models, which learn from vast text corpora, challenge the traditional view of language as a static set of terms and rules, instead proposing that language fundamentally consists of relational links between words [13]. This perspective aligns well with the core objective of KGs, which is to map out a network of relationships among entities. While LLM-based in-context learning (ICL) has demonstrated state-of-the-art performances in biomedical NLP tasks without expensive training or fine-tuning, their potential for end-to-end KG construction, particularly for MRPs, remains largely untapped and represents a promising frontier in the field of biomedical research [13]. Additionally, tools such as Bioinfo-Bench [14] are significant in evaluating the capabilities of LLMs in bioinformatics, indicating a promising direction for future research.

In this paper, we explored the capability of GPT-4 in the end-to-end construction of a context-aware relational graph to accurately delineate context-specific MRPs of m^6^A methylation within a given sentence. Our contributions are:

1. We proposed reguloGPT, GPT-4 driven ICL prompt, specifically designed for end-to-end joint NER, N-ary RE, and context identification, with an aim to accurately interpret context-specific MRPs that include both explicit and implicit regulations. We designed the baseline, few-shot, and Chain-of-Thought (CoT) prompts for reguloGPT.
2. We introduced a context-aware relational graph representation of regulatory interactions within MRPs of disease, tissue, and cell type (Fig. 1). This graph uniquely incorporates the context as part of the relational edges, thereby addressing and resolving the semantic contradictions of relations that often arise when contexts are not considered (Fig. 1). It also possesses the inherent regulatory hierarchy of MRPs (Fig. 1).
3. We annotated the context-aware relational graphs derived from 400 PubMed paper titles related to m^6^A MRPs and created a benchmark dataset. This dataset encompasses a diverse array of contexts, entities and relationships, highly valuable for systematic evaluation of reguloGPT.
4. We thoroughly evaluated the performance of the proposed prompts for predicting contexts, recognizing the entities, and extracting both explicit and implicit relationships. Our results demonstrated significant improvement over several existing algorithms.
5. To overcome the need for manual annotation in evaluating reguloGPT, we introduced a novel G-Eval scheme, which leverages CoT prompts to evaluate extracted context and relational graphs. We showed that there was a strong similarity between G-Eval scores and annotation-based evaluations.
6. We applied reguloGPT to PubMed titles between 2013-2023 related to m^6^A MRPs and constructed m^6^A-KG, a comprehensive KG of m^6^A MRPs. We demonstrated the utility of m^6^A-KG for representing m^6^A-mediated pathways and delineating mechanisms by which the m^6^A writer METTL3 regulates cancer-related phenotypes in breast cancer, lung cancer, and myeloid leukemia.

## II. METHODS

In this section, we outline ***reguloGPT***, a novel approach that leverages GPT-4 based ICL for the end-to-end extraction of MRPs from literature. The reguloGPT involves six modules, each meticulously designed to facilitate the construction of a context-aware KG from PubMed research publications, as illustrated in Fig. 2. The reguloGPT workflow begins with a dataset of publication titles extracted from PubMed. These titles are fed into reguloGPT, which utilizes a customized ICL prompt. The prompt is designed to capture N-ary molecular regulations and their biological context, reflecting the intricacies of MRPs. We will detail these processes in the subsequent sections, covering the generation, annotation, and normalization of the benchmark dataset for reguloGPT evaluation, evaluation criteria and methods, creation of a KG specific to m^6^A research domain, and the discovery of novel regulations.

**Fig. 2.**
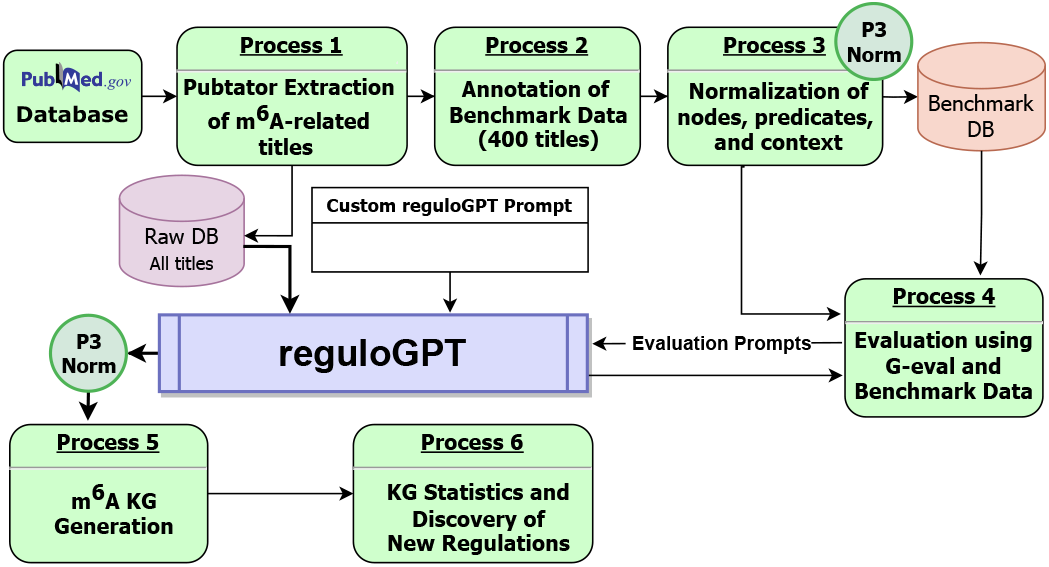
The overall process of developing reguloGPT including data collection, creation of a benchmark dataset, ICL prompt engineering, performance evaluation, context-aware m6A KG generation, and downstream analysis.

### A. In-Context Learning (ICL) Prompts for reguloGPT

ICL has gained prominence as an innovative method in LLMs, like GPT4, for zero-shot or few-shot predictions. To harness this potential, we developed three distinct prompts for reguloGPT including a baseline prompt that provides only definitions, a few-shot prompt enriched with a few examples that showcase the resultant context and N-ary relational graph, and a CoT prompt, which uses additional reasoning steps within each example, improving the underlying logic of the information extraction.

#### 1) Baseline prompt

Fig. 3A shows the framework of the baseline prompt, including: 1) Instruction, which presents the task objective of reguloGPT for GPT-4; 2) Definition, which defines the components in a context-aware relational graph, including node, edge, context, and inferred edge. Each edge includes two nodes and a predicate. This section also illustrates a collection of constraints for nodes and edge extraction; and 3) Output format. Following the prompt, we specify a target sentence from a PubMed paper that comprises a collection of molecular regulatory relationships. In this paper, we only use the title of a paper. In the definition, we also propose the inferred edge since many relationships in the sentences are logically derived but aren’t directly stated in the provided sentence. Take “METTL3-mediated m^6^A methylation of SPHK2 promotes gastric cancer progression by targeting KLF2” in Fig. 1 as an example, we can infer an edge for KLF2 promoting gastric cancer progression but the sentence does not explicitly mention this relationship.

**Fig. 3.**
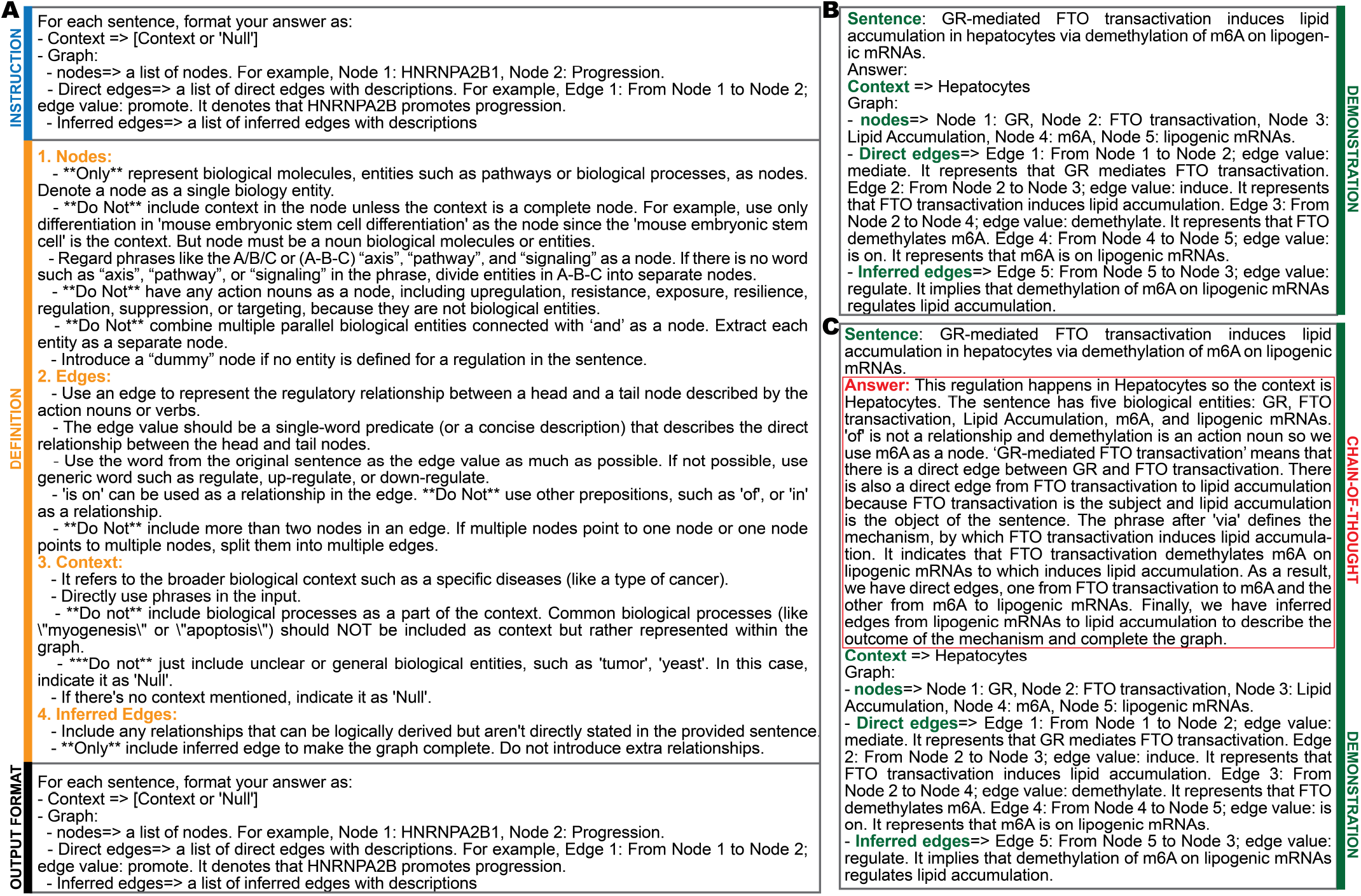
The reguloGPT prompts. (A) Baseline prompt including instruction, definition, and output format. (B) Demonstration in few-shot prompt. (C) Demonstration in CoT prompt.

#### 2) Few-shot prompt

The few-shot prompt consists of 1) instruction, 2) definition, 3) demonstration, and 4) output format. Different from the baseline prompt, the few-shot prompt includes an extra demonstration section after definition, which provides a few examples containing pairs of sentences and the biomedical graph extracted from sentences. A few examples help LLM have a better understanding of the task. We include 4 examples in our prompt and one of them is illustrated in Fig. 3B. Each example includes the target sentence and output (context, nodes, direct edges, and inferred edges). The output follows the requirement in the output format.

#### 3) Chain-of-Thoughts (CoT) prompt

CoT prompt has been shown [15] to encourage a complex and logical response from LLM, which in turn improves the task performance. In our CoT prompt, we add a series of intermediate reasoning steps as the chain of thought for each example in the demonstrations, as presented in the red box in Fig. 3C.

### B. Construction of datasets for performance benchmark and knowledge graph generation

The lack of context-dependent benchmark datasets for MRPs is a primary obstacle to the comprehensive assessment of our proposed reguloGPT. In addition, in the rapidly evolving field of molecular biology, the focused construction and annotation of benchmark datasets in the m^6^A domain hold significant scientific value. Concentrating on m^6^A, a relatively new area, allows for a detailed and nuanced understanding of this emerging field. This targeted approach not only circumvents the challenge of information overload inherent in broader research domains but also fosters the development of a specialized repository of knowledge. Such a repository is instrumental in accelerating research and catalyzing new discoveries in m^6^A-related studies. Moreover, the creation of a benchmark dataset within this niche is critical for model validation and refinement.

We extracted titles of publications involving m^6^A research as they represent the most concise description of context-specific molecular regulations. To this end, we searched PubMed, which is a large open database of online books, life science journals, and MEDLINE. We used PubTator [16] RESTful API with “m^6^A” as the query keyword to extract publications from PubMed between 2013 and 2023. Our selection criteria are defined as follows: we choose titles that are complete sentences and include references to multiple genes. This is crucial for mapping pathways that either lead from m^6^A to various genes/proteins or from these genes/proteins back to m^6^A.

#### 1) Annotation method for benchmark dataset

To facilitate the annotation of a benchmark dataset, we assembled five subject-matter-expert annotators with backgrounds in computer science and biomedicine to annotate 400 specially chosen titles, which contain MRPs from the m^6^A research paper title corpus. The annotation has three phases:

a. Practice annotation phase: We randomly selected 20 sentences as practice examples. Five annotators followed the descriptions that were provided in the prompts to identify the nodes, edges, and context. They discussed within themselves and came up with a consensus. Most importantly, they summarize the special cases for further annotation.
b. Group annotation phase: We used annotation guidelines summarized in the practice annotation phase to guide the group annotation. All sentences were divided into 5 shards and distributed to 5 annotators. After the first round of annotation, 5 annotators exchanged examples and completed the second round of annotation. In this case, each example was annotated by two annotators.
c. Adjudication phase: For titles that all annotators agreed on, their annotation will be final. For the others, the annotations were discussed within the group to reach an agreement.

#### 2) Annotation guidelines

In the practice annotation phase, basic guidelines were summarized. For each sentence, the annotation included context, nodes list, and edge lists. Each edge included two nodes and one predicate to connect the two nodes. Inferred edges were considered to be extra relationships and they were often accompanied by prepositions like “via”, “by”, and “through” in the sentence. The context should not be biological processes such as development, progression, etc. Co-reference was not committed and therefore for “m^6^A methyltransferase METTL3” only “METTL3” was extracted. In addition, some special cases were adopted: 1) Any complex mechanism like the A/B/C or (A-B-C) “axis”, “pathway”, and “signaling” is annotated as single node. If there is no word such as “axis”, “pathway”, or “signaling” in the phrase, divide entities in A-B-C into separate nodes; 2) A “dummy” node was introduced if no entity is defined for regulation in the sentence. For example in the Fig. 1, for the subject “Deficiency of METTL3” at the beginning of the sentence with PMID 33681207, we will construct a relationship as (dummy, induce deficiency of, METTL3). Finally, we normalized the extracted relationships into 31 ontological normalized predicates discussed in the next section.

### C. Normalization of nodes, predicates, and contexts

We used Gilda [17] and the Gene Ontology knowledgebase (GO) [18] to normalize nodes first. Subsequently, we performed manual normalization to ensure consistency among nodes that convey the same meaning. We further grouped the nodes into six categories: m^6^A, m^6^A writers/erasers/readers (WERs), genes/proteins, GO/pathways, and other.

For the predicate normalization, we followed the Ontological predicate definitions in SemRep [4]. Semrep provides 30 predicate types including HIGHER THAN, LOWER THAN, AFFECTS, STIMULATES, AUGMENTS, INTERACTS WITH, INHIBITS, DISRUPTS, PREVENTS, CAUSES, DIAGNOSES, CONVERTS TO, COEXISTS WITH, COMPLICATES, ISA, TREATS, PRODUCES, LOCATES, PRECEDES, MANIFESTS, METHODS, OCCURS IN, PART OF, COMPARED WITH, SAME AS, ASSOCIATED WITH, USES, ADMINISTERED TO, PROCESS OF, PREDISPOSES. We added an extra predicate type, MAINTAINS (keep in an existing state) to have 31 types in total. For relationship normalization, we applied GPT-4 to perform an initial normalization, followed by a manual evaluation to correct inconsistencies. We also applied the same normalization method to the context as we do for nodes. We further systematically normalized the contexts associated with The Cancer Genome Atlas (TCGA) cancer types [19].

### D. Construction of the m^6^A knowledge graph

In addition to the benchmark dataset of 400 titles, our study further extracted 968 titles that include descriptions of MRPs from the titles extracted by PubTator. These additional titles were subject to our reguloGPT CoT prompt to extract the context and relation graphs, thus broadening the scope of our analysis and enriching the dataset under consideration. Normalization was applied to standardize the extracted nodes, edges, and contexts. We integrated these normalized relational graphs with those from our benchmark dataset by joining common nodes and edges to construct m^6^A-KG, a comprehensive KG of m^6^A functions in diverse contexts. This KG includes nodes connected with edges that define the normalized predicates. A unique feature of m^6^A-KG is that each edge also includes a set of associated contexts extracted from the same titles as the edge to inform the context under which the regulation defined by the edge occurs. The edge also incorporates the unnormalized edge value and PubMed Identifier (PMID) of the associated titles. Unnormalized edge and PMID provide a mechanism to trace back to the original title and associated paper for reference. We used Neo4j [20] to visualize and manipulate our KG.

### E. Evaluation metrics and criteria

#### 1) Evaluation with the benchmark dataset

We used the benchmark dataset to evaluate the performance of reguloGPT across different prompt designs. We adopted accuracy as the metric for context prediction and recall, precision and F1 score for nodes and edges evaluation. The criteria to evaluate the predicted nodes and edges are listed below:

- **True positive**: This is achieved when GPT-4 prediction nodes align with the benchmark annotation. A match is also considered if the output context or node contains most of the ground truth information. For edge evaluation, the criteria for two nodes are similar, and the normalized prediction must completely align with the result in the benchmark dataset.
- **False positive**: Incorrectly extracted nodes or edges are marked as false positives. In edge evaluation, a false positive occurs if the predicted nodes match but the predicate is incorrect or not extracted.
- **False negative**: Any ground truth nodes and edges without a corresponding matching prediction are false negatives.

*2) G-Eval scheme for annotation-free assessment of reguloGPT:*

The assessment of context-aware KG construction poses challenges and manual annotation is labor-intensive and costly. Recent research proposes leveraging LLMs directly as evaluators for reference-free Natural Language Generation, as indicated by [21] in GPTScore. They utilize LLMs to evaluate candidate outputs, assigning scores based on generation probability without referencing any target. [22] demonstrate that GPT-4 can assess the quality of generated texts in coherence, consistency, fluency, and relevance compared to ground truth in a form-filling paradigm. However, existing studies have primarily focused on sentence-level evaluation, leaving the performance of LLMs in graph generation evaluation largely unexplored.

To address this challenge, we proposed a novel framework, GPT-4-evaluation (G-Eval), which employs GPT-4 and a form-filling paradigm to evaluate the quality of output at the sentence level. We experimented with two tasks, namely, 1) context evaluation and 2) graph evaluation. For context evaluation, GPT-4 gave a score to each context in a sentence, while for graph evaluation, GPT-4 gave a score to all edges extracted from a sentence. The evaluation prompts of both context evaluation and graph evaluation included four parts: 1) Introduction, 2) Definition, which denotes the concept of context in the context evaluation, or the concepts of nodes and edges in the graph evaluation; 3) Evaluation Steps; and 4) Output Format.

The concepts of contexts, nodes, and edges are the same as those defined in reguloGPT prompts (Fig. 3A). The evaluation steps were generated by GPT-4 based on the introduction and definition. The range of the score was 1-5, and we repeated the evaluation five times to obtain the average score [22]. In the output format, we added a test sentence and predicted context in the context evaluation or corresponding edge (two nodes and a predicate) list in the graph evaluation. It should be mentioned that we used unnormalized context and edges in the output. Fig. 4 shows the framework of G-Eval for the context evaluation and graph evaluation.

**Fig. 4.**
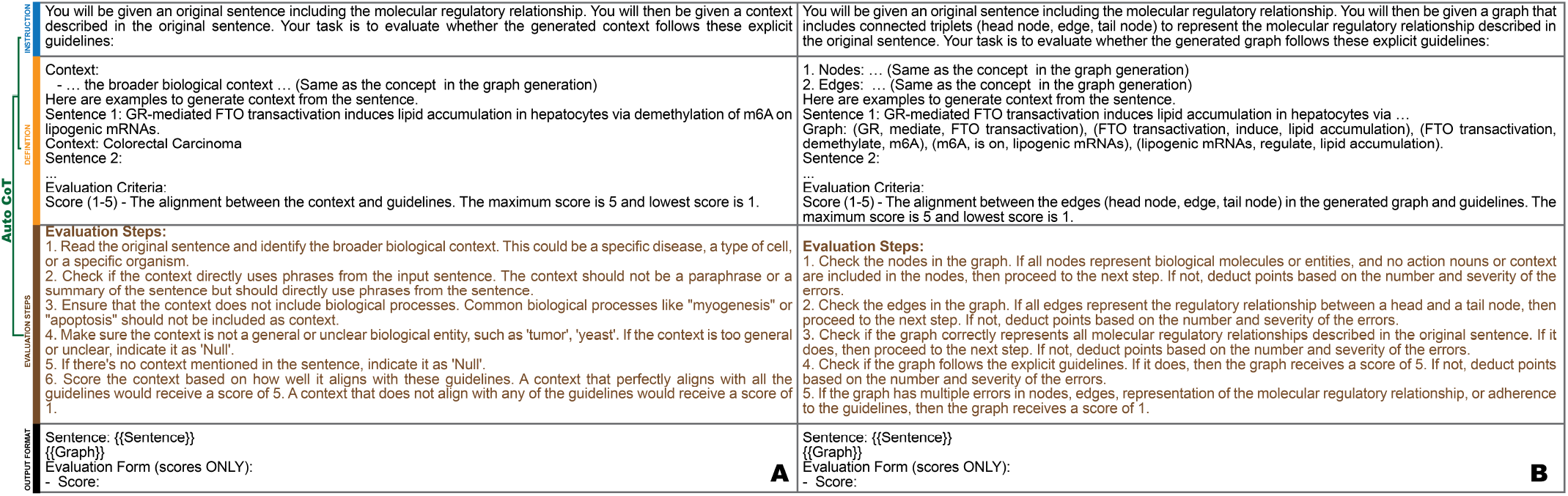
The G-Eval prompts for (A) context evaluation and (B) graph evaluation. the Evaluation Steps were generated by GPT-4 based on our Instructions and Definitions. Then, they evaluate the context or graph added in the Output Format in a form-filling fashion.

## III. Results

### A. Annotation of the benchmark dataset

We annotated the context-aware graphs for a selection of 400 titles, specifically chosen from m^6^A research papers. We were able to deduce the context-specific information from 344 titles. The annotated dataset includes the extracted 1558 nodes and 1485 edges with 1312 unique nodes and 152 unique edges, or an average of 3.72 entity-relations extracted per title. Further normalization resulted in a total of 1241 unique nodes and 62 unique edges. Also, 165 of the nodes were categorized as in the Genes/Proteins group, 172 as GO/Pathway, 9 as Readers, 8 as Writers, 2 as Erasers, and 956 as Other. Moreover, we were able to extract 24 different TCGA cancer types from the normalized contexts in the benchmark dataset.

### B. reguloGPT significantly outperforms existing algorithms on the benchmark dataset

We first evaluate reguloGPT’s performance on the benchmark datasets against human annotation. To evaluate the effectiveness of reguloGPT, we selected two established algorithms as baselines: REACH [5] and EIDOS [6]. Both algorithms are integral components of the INDRA [3] framework and are specifically designed for extracting interactions from scientific research papers. To conduct a comprehensive comparison, we tested these baseline algorithms using the benchmark dataset. Note that neither baseline algorithms were designed to extract contexts.

Table I details the performance of various prompting strategies used in reguloGPT development (baseline, few-shot, and CoT prompts) compared to REACH and EIDOS, as measured against the human-annotated benchmark dataset. The metrics used for this comparison include Recall (Re), Precision (Pr), and F1 score for both node and edge evaluations, alongside Accuracy for context evaluation. Because REACH and EIDOS do not output context information, hence context evaluation results (accuracy) for these algorithms are absent in the comparison.

**TABLE 1.**
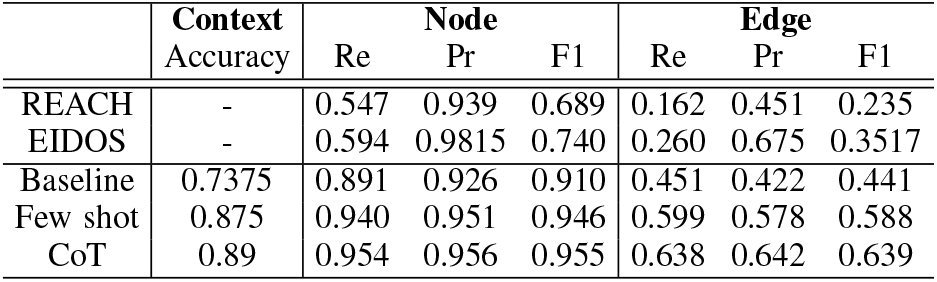
Results comparison of different prompts with existing algorithms using the benchmark dataset. Here, Re = Recall, Pr = Precision, and F1 = F1 score.

Overall, reguloGPT’s ICL strategies have demonstrated remarkable superiority over REACH and EIDOS. Among reguloGPT prompts, CoT emerged as the most effective, achieving an impressive accuracy of 0.89 for context detection and F1 scores of 0.955 for node prediction and 0.636 for edge extraction. The relatively lower performance on edge prediction underscores the inherent complexity in accurately extracting complex N-ary relationships. However, when compared to EIDOS, the CoT prompt showed substantial improvement of 22%, 29%, and 81.5% improvement in context accuracy and node and edge F1 scores, respectively. These enhancements underscore reguloGPT’s overall superior capabilities in extracting knowledge of MRPs. The marked improvement can be attributed to the end-to-end strategy and, likely, the advanced capabilities of GPT-4.

The improvement of the extraction capabilities is evident in the title *“The m^6^A methyltransferase METTL3 promotes osteosarcoma progression by regulating the m^6^A level of LEF1”* (PMID: 31253399). As noted in section III-A, the benchmark annotations for this title include four triplets under the context of ‘osteosarcoma’. However, REACH only identified (METTL3, STIMULATES, level of LEF1). Similarly, EIDOS extracted only one triplet (m^6^A methyltransferase METTL3, STIMULATES, osteosarcoma progression). In contrast, all three of the reguloGPT prompts were able to successfully extract the 3 direct and 1 inferred edge relationship between the correct entities with the correct context of osteosarcoma.

In another example, *“eIF3i promotes colorectal cancer cell survival via augmenting PHGDH translation”* (PMID: 37611825), reguloGPT identified three triplets with two direct and one inferred edge. In contrast, REACH extracted only one triplet (eIF3i, STIMULATES, cell survival) while EIDOS extracted two triplets, including (eIF3i, AUGMENTS, PHGDH translation) and (eIF3i, STIMULATES, colorectal cancer cell survival). However, reguloGPT was able to additionally extract the inferred edge relation (PHGDH, STIMULATES, survival) and the context as ‘colorectal cancer cell’.

Due to additional demonstrations in the prompts as illustrated in Fig. 3C, the CoT prompt leads to Context accuracy at 89%, followed by the few-shot prompt at 87.5%, and the baseline prompt at 73.75%. In extracted Node evaluation, the CoT prompt again demonstrates superior performance, achieving the highest scores across Recall (95.4%), Precision (95.6%), and F1 (95.5%) followed by a few-shot prompt that surpasses the other methods. For Graph evaluation, CoT leads with a Recall of 63.8%, Precision of 64.2%, and an F1 score of 63.9%. The few-shot prompt closely follows, significantly outperforming the baseline prompt, EIDOS, and REACH algorithms.

To be precise, the advanced prompt technique makes the output of GPT-4 align with our requirements. Although we ask the GPT-4 to introduce a dummy node in the prompt, the output of the baseline prompt ignores this guideline. By adding one example in a demonstration with a similar case, the few-shot prompt can follow this requirement. However, this alignment is not stable. In the paper “*Suppression of m^6^A reader Ythdf2 promotes hematopoietic stem cell expansion*” (PMID: 30065315), the few-shot prompt neglects this condition, but the CoT prompt can maintain alignment as well. A similar issue happened in the “*Silencing METTL3 inhibits the proliferation and invasion of osteosarcoma by regulating ATAD2*” (PMID: 32044716) and the few-shot prompt fails to introduce a dummy node.

### C. G-Eval assessment is consistent with manual evaluations

We next investigated the G-Eval evaluations of predictions by the three reguloGPT prompts on the 400 titles in the benchmark dataset and assessed the extent to which the G-Eval evaluations are consistent with the evaluations against human annotations. We have 400 scores in context evaluation and 400 scores in graph evaluation. Examining the averaged scores across the 400 titles (Table. I) revealed a consistent trend with the annotation evaluation in Table. II where the CoT prompt exhibited the best performance, followed by the few-shot and baseline prompts.

**TABLE 2.**
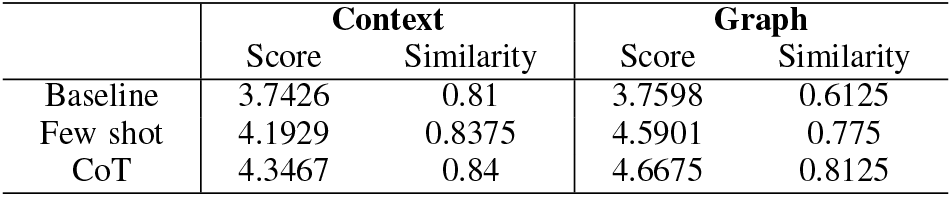
G-Eval results. The range of scores is 1 - 5. The similarity denotes the Rand similarity coefficient between the G-Eval and the human annotation evaluations of reguloGPT’s prediction on the benchmark dataset at the sentence level.

|  | Context |  | Graph |  |
| --- | --- | --- | --- | --- |
|  | Score | Similarity | Score | Similarity |
| Baseline | 3.7426 | 0.81 | 3.7598 | 0.6125 |
| Few shot | 4.1929 | 0.8375 | 4.5901 | 0.775 |
| CoT | 4.3467 | 0.84 | 4.6675 | 0.8125 |

To further validate the effectiveness of our G-Eval strategy, we analyzed the similarity between the annotation evaluation and G-Eval scores for each sentence. Since the annotation evaluation for each sentence is binary, i.e., correct or incorrect, we first binarized G-Eval scores using a threshold score of 3.

The threshold was chosen based on the score distribution (1-5). Additionally, G-Eval conducts the graph evaluation, whereas the annotation evaluations are assessed for nodes and edges. To make them comparable, we generated a graph-level annotated evaluation such that a sentence was deemed correct if more than 50% of the edges in the sentence were correctly predicted. We did not consider node prediction because their F1 scores are high as shown in Table. I. To compare the similarity between the G-Eval and annotation evaluations, we computed the Rand matching coefficient for each title. These results are detailed in Table. II. They demonstrate high similarities between the two evaluations, especially for reguloGPT, where the Rand similarities reach 0.84 for context prediction and 0.8125 for graph prediction. These results suggest that G-Eval is a promising annotation-free method for evaluating reguloGPT.

## IV. M^6^ A-KG, A CONTEXT-AWARE KG OF M^6^ A regulatory functions

m^6^A is the predominant mRNA modification in mammalian cells, present in over 40% of transcripts. The dynamic m^6^A regulation involves various RNA binding proteins (RPBs) including writers (METTL3 & METTL14), which add methyl groups, erasers (ALKBH516 & FTO2) to remove it, and readers, (e.g. YTH proteins), which bind to m^6^A sites to decode the regulatory signals for mediating gene expression. It achieves this by regulating mRNA stability, splicing, mRNA export, and translation efficiency. Additionally, it influences cancer development and progression significantly by modulating mRNA stability and splicing. Despite growing interest, the roles of m^6^A and its writers, erasers, and readers in cancer through gene expression alterations are not fully understood. We demonstrate the utility of reguloGPT to create a detailed representation of the m^6^A-associated molecular regulatory pathways.

### A. Construction of m^6^A-KG with reguloGPT

We applied reguloGPT to 968 unannotated titles, resulting in the extraction of context-aware relational graphs that depict functions related to m^6^A in diverse contexts. After normalizing the nodes, edges, and contexts, we synthesize these relational graphs and annotated graphs from the benchmark dataset into a comprehensive m^6^A knowledge graph (m^6^A-KG), denoting molecular regulatory pathways linked to m^6^A. The constructed m^6^A-KG comprises 2,397 nodes, 4,694 edges, and 478 unique contexts, with each edge encompassing an average of 1.06 contexts. The node degree, calculated by aggregating in-degrees and out-degrees akin to undirected graphs, follows a powerlaw distribution, with 96.2% of nodes having less than 10 degrees and only 9 nodes possessing *>*100 degrees. Notably, node “m^6^A” emerges as the most connected, with a degree of 827, highlighting its centrality in the network. The top nodes by degree include key m^6^A writers like METTL3 (436) and METTL14 (122), erasers such as ALKBH5 (166) and FTO (222), and readers like YTHDF2 (127) and YTHDF1 (109). This underscores their vital roles in the regulatory functions of m^6^A. Additionally, nodes representing cell proliferation (104) and neoplasm metastasis (93) also have high degrees, indicating m^6^A’s significant influence on these tumor-related phenotypes.

### B. The structure of m^6^A-KG reflects the architecture of molecular regulatory pathways

To examine if the structure of the m^6^A-KG reflects a typical architecture of MRPs, we categorized the nodes into six groups: **m**6**A**, m^6^A writers/erasers/readers (**WERs**), **GO/pathway, genes/proteins**, and **other**. Analysis of the outgoing edge percentage of a node (outdegree rate) within each group revealed a hierarchical structure aligned with that of a molecular pathway. Specifically, m^6^A WERs and m^6^A have a median 0.85 and 0.77 outdegree rate, respectively, suggesting that they occupy upstream positions (Fig. 5A) and re-affirming their role as key regulators. Also, **genes/proteins** nodes (0.05 median outdegree rate) are intermediate nodes, which bridge the upstream regulators with the downstream **GO/Pathway** nodes (0.03 median outdegree rate) (Fig. 5A). The **other** nodes exhibited three subgroups, with two (**other-L** and **other-H**) characterized by a median outdegree rate of either 0 or 1 (Fig. 5A), indicating their positions at extreme ends of the pathway. Close inspection revealed that **other-L** nodes define disease phenotypes or outcomes, naturally at the bottom of pathways, while **other-H** nodes include chemical or environmental stimuli and are expected to be upstream of pathways (Fig. 5B). The emergent structure of the m^6^A-KG, with various stimuli on the top followed by clear upstream m^6^A regulators, gene/protein interactions, and downstream phenotype outcomes, exhibits the hallmarks of an MRP.

**Fig. 5.**
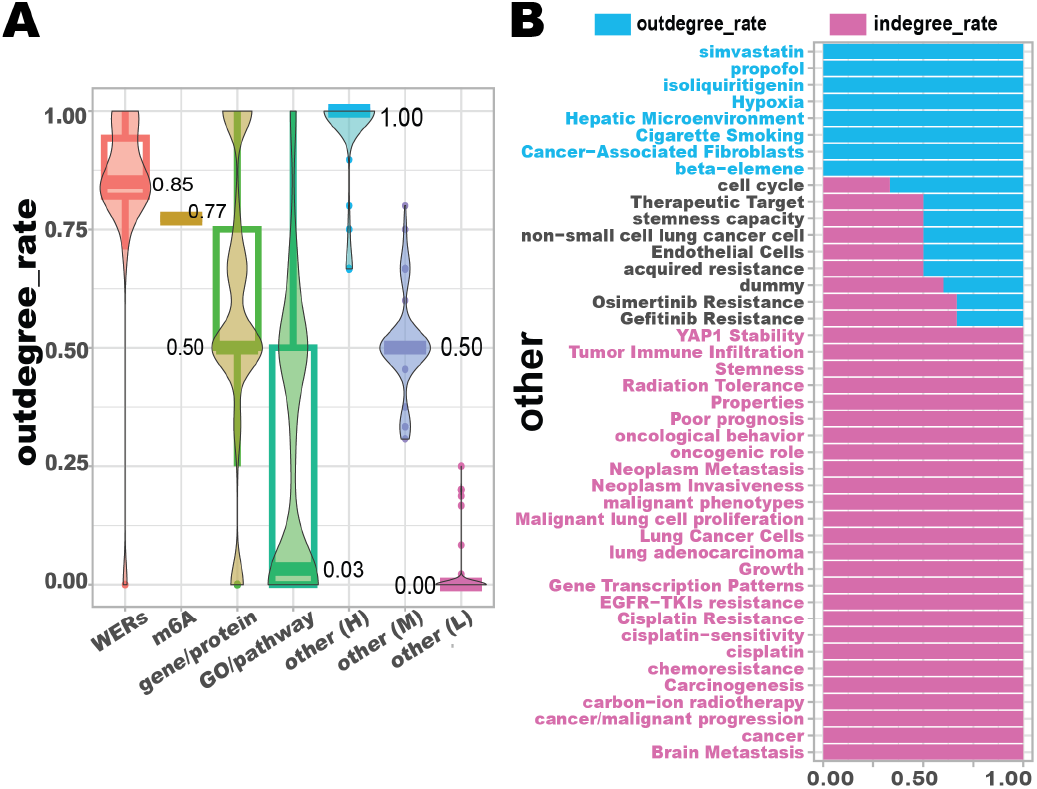
(A) Outdegree rate of nodes in different categories. (B) Outdegree and indegree rates of **other** category nodes within lung cancer-specific KG.

**Fig. 6.**
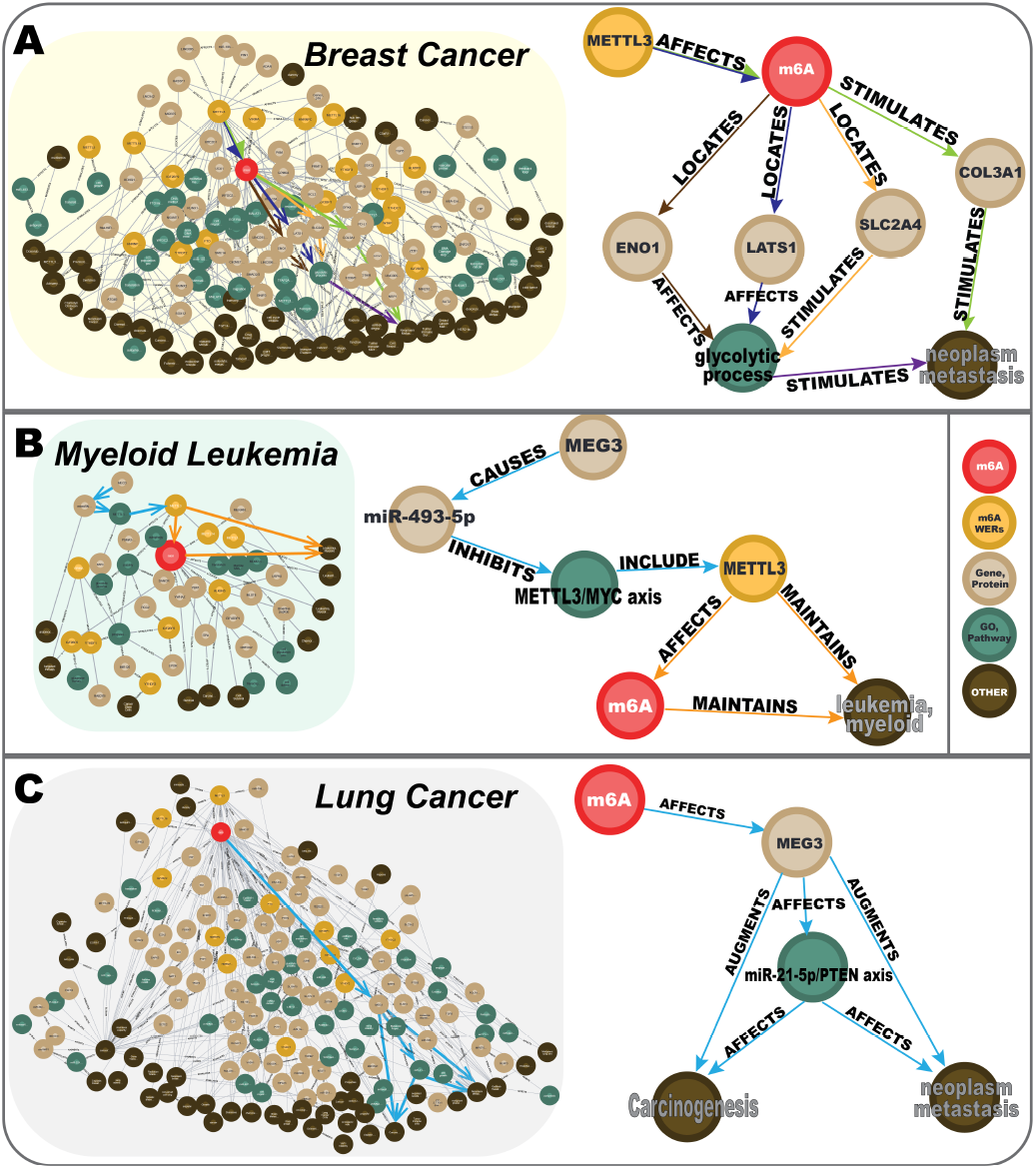
Cancer-type specific KG of (A) Breast cancer, (B) Myeloid leukemia, and (C) Lung cancer. Extracted pathways are shown to the left. Edge colors are associated with the supporting titles.

### C. The m^6^A-KG reveals distinct mechanisms of m^6^A functions across various cancer types

We next investigated m^6^A’s role in various cancers, leveraging the m^6^A-KG’s integration of contexts and PMIDs into edges. This feature enabled us to dissect functions specific to certain cancers and to identify those common across multiple types. The m^6^A-KG contexts included 24 TCGA cancer types with 2,366 edges pertaining to individual cancer types. Remarkably, one edge representing “METTL3 AFFECTS m^6^A” is universally presented across all 24 TCGA cancer types examined, signifying METTL3’s ubiquitous influence. Additionally, three edges spanning 10 cancer types involve the relationships “METTL14 AFFECTS m^6^A”, “ALKBH5 AFFECTS m^6^A”, and “METTL3 STIMULATES progression”, highlighting the central role of the m^6^A writers METTL3, METTL14, and the eraser ALKBH5 in multiple cancers.

To gain insights into cancer-specific m^6^A-mediated functions, we extracted cancer-specific KGs for breast cancer, lung cancer, and myeloid leukemia. These sub-KGs presented clear hierarchies of MRPs, with m^6^A regulators at the top and disease phenotype nodes at the downstream. METTL3’s widespread association across cancers prompted further examination of pathways centering on this regulator. We focused on pathways supported by edges spanning multiple titles because they could reveal novel functions. The breast cancer sub-KG delineates a complex dual-pathway mechanism, with evidence from five titles (PMID: 32766145, 36069931, 36609396, 34312368, 35319018), suggesting METTL3’s involvement in tumor metastasis through two distinct routes: regulation of COL3A1, crucial for extracellular matrix structure, and alteration of cancer cell metabolism via the glycolytic pathway. This duality suggests that therapeutic targeting METTL3 could simultaneously disrupt key structural and metabolic routes essential to cancer metastasis, offering a promising avenue for multifaceted therapeutic intervention. Moreover, cancer-dependent regulations of MEG3, a tumor suppressor gene, were revealed in lung and leukemia sub-KGs. The leukemia sub-KG indicates that MEG3 modulates miR-493-5p to suppress myeloid leukemia by inhibiting METTL3-mediated m^6^A methylation (PMID: 35761379, 29186125). Conversely, in lung cancer, METTL3 methylates MEG3, which facilitates carcinogenesis and neoplasm metastasis (PMID: 37308993). These distinct regulatory mechanisms were corroborated through a detailed examination of the literature associated with the extracted pathways, validating the m^6^A-KG’s utility in uncovering new functional insights.

## V. Conclusion

In this study, we introduced reguloGPT, a novel application of GPT-4 for the end-to-end construction of KGs in the realm of MRPs. We developed ICL prompting strategies to extract context-aware relational graphs depicting interactions with MRPs. We thoroughly evaluated reguloGPT’s efficacy against a human-annotated benchmark database comprising 400 titles and demonstrated significant improvements over existing algorithms. We also found a good similarity between manual evaluation and our proposed annotation-free G-Eval. We successfully applied reguloGPT to create a comprehensive and detailed m^6^A-KG. This KG included an extensive network of 2,397 nodes and 4,694 edges, providing a rich map of m^6^A regulatory functions. A notable feature of m^6^A-KG is its unique context-aware edges, which incorporate associated contexts and PubMed IDs. This design not only allow us to understand context-specific regulations but also improves traceability and verification of the data. The m^6^A-KG revealed distinct mechanisms of m^6^A functions across various cancer types, facilitating a deeper understanding of the role of m^6^A in cancer, opening avenues for targeted cancer research and therapy development. The hierarchical structure of the m^6^A-KG mirrors the architecture of MRPs, revealing a more intuitive understanding of the complex interactions and roles within these pathways. Future studies will explore a more systematic G-Eval assessment and relationship extraction, along with improved normalization schemes for edges and contexts. A systematic and effective approach to elucidate novel regulatory functions from the KG will be further developed.

## VII. Acknowledgments

This work is supported in part by funds from the National Institutes of Health (U01CA279618, 1SC3GM136594-02, R01CA124332, U24CA269436, R00CA248944, 3R00CA248944-04S1, R03OD036494, P30CA047904, and P30DK120531) and the Leukemia Research Foundation.

